# Exploring Bioelectricity with Ace2N-mNeon during Zebrafish Embryogenesis

**DOI:** 10.1101/2024.12.12.628143

**Authors:** ZhenZhen Wu, Rui Oliveira Silva, Ruya Houssein, Fabiola Marques Trujillo, Jordan Gotti, Srividya Ganapathy, Zhenyu Gao, Daan Brinks

## Abstract

Bioelectricity is a fundamental biophysical phenomenon present in all cells, playing a crucial role in embryogenesis by regulating processes such as neuronal signaling, pattern formation, and cancer suppression. Precise monitoring of bioelectric signals and their dynamic changes throughout development is vital for advancing our understanding of higher organisms. However, the lack of suitable techniques for mapping bioelectric signals during early development has greatly limited our ability to interpret these mechanisms. To address this challenge, we developed an Ace2N-mNeon expression library in zebrafish, which exhibits membrane localization from 4 hours post-fertilization to at least 5 days post-fertilization and broad expression across multiple cell types throughout development. We validated the use of this library for studying bioelectric changes via voltage imaging to record signals in neurons and cardiomyocytes at different development stages. Through this approach, we found evidence of synchronized neuronal activity during early embryogenesis and observed faster voltage dynamics in cardiomyocytes as development progressed. Our results show that the Ace2N-mNeon library is a valuable tool for developmental bioelectric studies supporting advanced techniques such as voltage imaging and fluorescence lifetime imaging (FLIM). These methods enable non-invasive, dynamic monitoring of bioelectric signals across diverse cell types throughout development, significantly surpassing the capabilities of current electrophysiological techniques.

## 1. Introduction

Embryogenesis is a crucial phase in the development of higher organisms, and its study is essential for understanding the complex processes of growth and cellular differentiation. Among the various factors influencing this process, bioelectricity, defined as the voltage changes and ionic currents across cellular membranes^1^, is thought to be a significant regulator^2–5^. These bioelectric signals span across a broad range of timescales, from rapid action potentials (APs) changes within milliseconds in neurons or muscle cells^5^, to slower, long-term changes in resting potentials (RPs) lasting from minutes to days in all cell types^3,6^. Understanding these dynamic bioelectric signals is important for decoding their roles in cellular behavior during development^3^.

Neuronal APs, also known as spikes, encode and transmit information to support functions such as sensory perception, motor control, and cognition^7^. Similarly, cardiac APs are important markers of heart function, originating from the interplay of membrane proteins that regulate muscle contraction^8^. In contrast, slower changes in RP, often described as endogenous electric fields or developmental bioelectricity^9^, which emerges as early as the two-cell stage in vertebrates^10^, regulate diverse processes such as cell proliferation^11,12^, differentiation^12^, migration^13^, polarization^14^, and apoptosis^4^. Moreover, developmental bioelectricity orchestrates cellular behavior by coordination with other physiological signals, thereby establishing positional information and anatomical identity^14^. It also plays important roles in wound healing^13^, appendage regeneration^15^, organ symmetry^15^, and cancer suppression^6,16–19^.

To advance our understanding of bioelectric signals during embryogenesis, it is crucial to simultaneously monitor both rapid APs and slow RPs in multiple cells within live organisms during early development. However, this remains technically challenging due to the spatial and temporal constraints of current electrophysiological techniques^20–22^. Recent advances in genetically encoded voltage indicators (GEVIs) have significantly improved our ability to study these processes^23–26^, enabling non-invasive, real-time visualization of membrane potential (Vm) changes through fluorescence^24,27^. While GEVIs have been widely used to study fast-spiking neuronal activities and cardiac dynamics, their application in early developmental stages remains limited^28–30^. Moreover, voltage imaging traditionally relies on correlating fluorescence changes to voltage changes, instead of measuring absolute RPs of cells. Additionally, while developmental bioelectricity studies often link bioelectric signals to physiological events^2,5,14,31,32^, off-target and compensatory effects from manipulating bioelectricity can lead to unpredictable results^2^. Therefore, accurately monitoring the spatial and temporal changes of Vm is crucial for understanding the role of bioelectricity during development. However, technical limitations have hindered the spatial mapping of absolute Vm over extended periods^2,33^. Recent studies using approaches such as fluorescence lifetime imaging (FLIM) have shown promise for estimating RP changes in live organisms^33–35^, providing a new way for exploring long-term bioelectric dynamics during early development. Thus, with the combination of voltage imaging and FLIM, researchers are gradually provided with the tools to analyze both APs and RPs during early development.

Zebrafish have become a preferred model for embryonic development research^36,37^ due to their small size, high embryo throughput, and genetic similarity to humans^37,38^. Additionally, their optical transparency further facilitates advanced imaging techiniques^39^, making them ideal for in vivo voltage imaging to study developmental bioelectricity. Several studies have successfully used GEVIs to image bioelectric activities in zebrafish brains^40–42^ and heart^29,43–45^, with some also exploring their application during early development stages^46,47^. These studies highlight the potential of combining zebrafish with GEVIs for studying bioelectricity in embryogenesis. Among the various available GEVIs, Ace2N-mNeon shows remarkable performance^48^, making it suitable not only for monitoring APs but also for mapping long-term RP changes through FLIM^34^.

In this study, we explored the potential of Ace2N-mNeon for investigating bioelectric signals during zebrafish embryogenesis. Initially, we characterized its ubiquitous expression patterns throughout development. We then developed an Ace2N-mNeon expression library using cell-specific promoters, enabling targeted studies of bioelectricity in distinct cell populations. To demonstrate the feasibility of these constructs for studying Vm, we tracked spontaneous activity in multiple neuronal populations during early brain development and found synchronization in neuronal populations during primary but not secondary neurogenesis. Additionally, we mapped cardiac AP propagation in embryonic zebrafish hearts, revealing chamber-specific transitions and faster voltage dynamics with distinct waveform patterns as development progressed. Our findings highlight the potential of Ace2N-mNeon for studying bioelectric signals in embryogenesis, providing novel insights into their roles in developmental processes.

## 2. Results

### 2.1 Ace2N-mNeon Predominantly Localizes to the Plasma Membrane Across Diverse Cell Types in *Danio rerio* During Development

To study bioelectricity during development, we explored strategies to express Ace2N-mNeon at different developmental stages. First, to observe the earliest expression, Ace2N-mNeon mRNA was injected into one-cell-stage fertilized eggs at the cytoplasmic region (Fig. 1a). Membrane-localized expression of Ace2N-mNeon was first detected at 2.75 hours post-fertilization (hpf) in positive embryos, corresponding to the onset of the blastula stage (Fig. 1b).

**Fig. 1.**
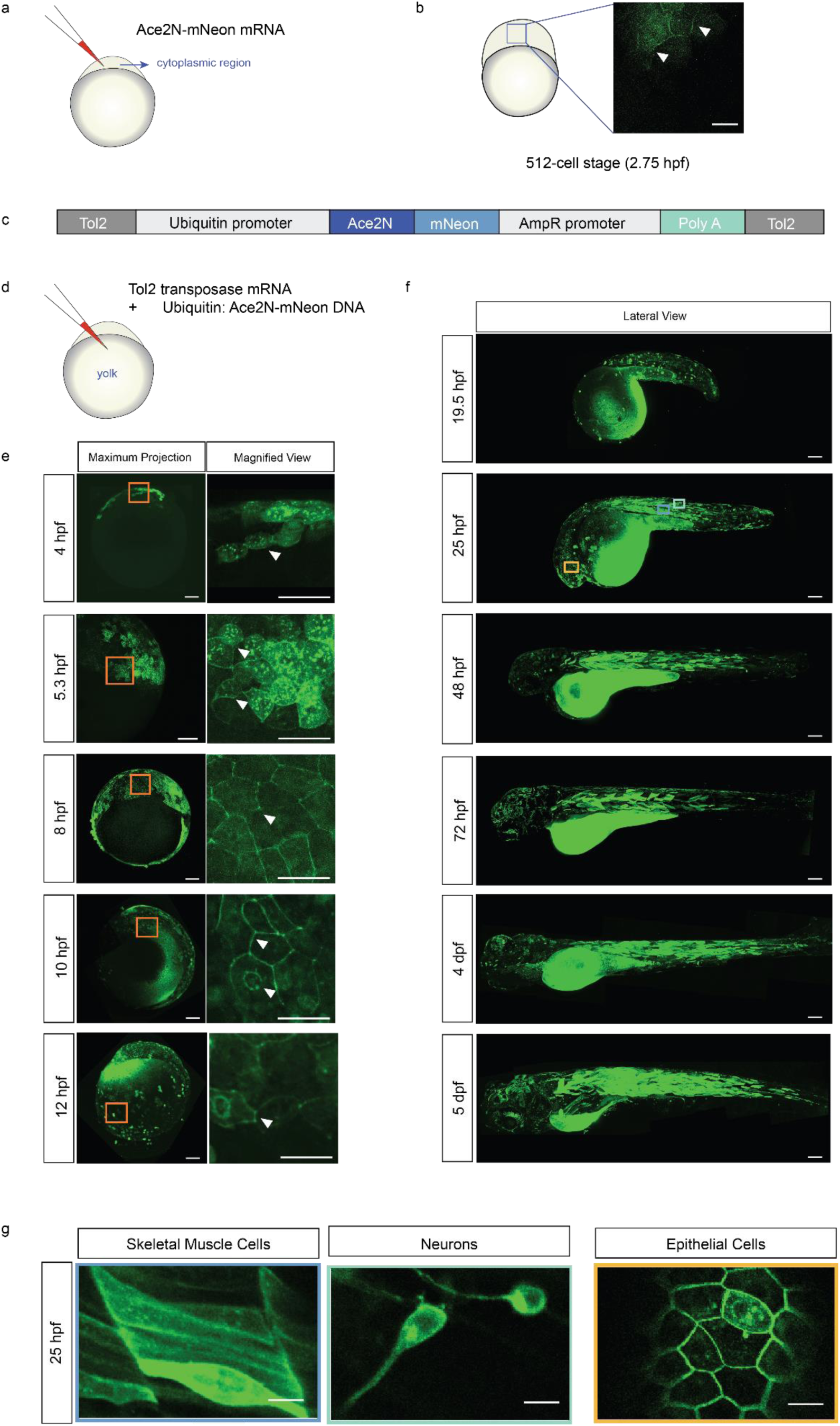
Expression of Ace2N-mNeon in Various Cell Types Throughout Development. (a) Schematic of microinjection of Ace2N-mNeon mRNA into the cytoplasm of a 1-cell stage embryo. (b) Single z-plane confocal images of Ace2N-mNeon mRNA expression at the 512-cell stage. Membrane localization can be observed starting at 2.75 hpf (white arrows). Scale bar = 20 μm. (c) Diagram of plasmid constructs for Ace2N-mNeon driven by the ubiquitin (ubi) promoter using the Tol2 transposon system. (d) Schematic of co-injection of Ace2N-mNeon-ubiquitin plasmid and Tol2 transposase mRNA into the yolk at the 1-cell stage. (e-f) Maximum projection images of zebrafish under the ubi promoter from sphere stage (4 hpf) to 5 dpf. Ace2N-mNeon is widely expressed and localized to membranes across various cell types. (e) Early expression from 4 hpf to 12 hpf. Left: Maximum projection, scale bar = 100 μm. Right: Magnified view showing membrane localization, scale bar = 50 μm. (f) Lateral view of Ace2N-mNeon expression in multiple cell types from 1 dpf to 5 dpf. Scale bar = 100 μm. (g) Single z-plane confocal images of Ace2N-mNeon expression at 25 hpf. Distinct cell types can be identified based on morphology, including skeletal muscle cells, neurons, and epithelial cells. Scale bar = 20 μm. hpf: hours post-fertilization; dpf: days post-fertilization.

To further investigate Ace2N-mNeon expression throughout zebrafish development, we used the Tol2 transposon system, a widely used tool in molecular biology enabling stable integration of transgenes into the host genome via a "cut-and-paste" mechanism. Upon microinjection, the transposase protein, translated from the injected mRNA, mediates the integration of the transgene into the zebrafish genome, leading to higher and more widespread expression compared to episomal plasmid maintenance. This method ensures greater consistency and stability of transgene expression across different cells and embryos, enhancing the reliability of the observed phenotypic outcomes^49^.

First, we studied the broad expression of Ace2N-mNeon using this system under the ubiquitin (ubi) promoter. Ubi can drive robust and ubiquitous transgene expression throughout zebrafish development. Previous studies have shown the introduction of EGFP across various cell types from 4 hpf, which continues throughout development with variable expression levels across different cell types^50^. We constructed a pTol2pA2_ubi_Ace2N- mNeon plasmid using a Tol2 transposon system (Fig. 1c) and co-injected it with Tol2 transposase mRNA into the yolk of one-cell-stage embryos (Fig. 1d).

Ace2N-mNeon showed broad expression in multiple cell types, which increased at later developmental stages, consistent with the expected timing of gene expression in cell differentiation^51,52^ (Fig. 1e, f). Positive embryos showed Ace2N-mNeon expression starting from the sphere stage (4 hpf) through confocal microscopy. As the embryos progressed into the gastrula stage, from 50% epiboly (5.3 hpf) to the formation of the embryonic shield (10 hpf), Ace2N-mNeon expression persisted in newly differentiated cells, with strong fluorescence at the cell membrane (Fig. 1e, 4-8 hpf). From 10 hpf, as segmentation began and somites formed (Fig. 1e, 10-12 hpf), the embryo developed a more defined body shape by the 21-somite stage (19.5 hpf) various cell types became distinguishable (Fig. 1f, 19.5 hpf). By the prim-6 stage (25 hpf), the embryo exhibited a recognizable body plan, including head, tail, and clear segmentation (Fig. 1f, 25 hpf). At this stage, multiple cell types were identifiable by morphology, including skeletal muscle, neural, and epithelial cells, with strong expression in the cell membrane (Fig. 1g). This expression can be observed throughout development, at least until 5 days post-fertilization (dpf). More detailed expression images from 1 dpf to 5 dpf can be found in Supplementary Fig. 1.

In all injected fish, we observed strong fluorescence in the yolk-sac throughout development as well, regardless of the construct used. To understand if this was an underlying artifact from the injection strategy, we imaged control non-injected zebrafish at the same development stages (Supplementary Fig. 2). Interestingly, control zebrafish also presented fluorescence in the yolk-sac. Thus, we disregarded this yolk-sac autofluorescence for the expression assessments. Our findings demonstrate the stable and widespread expression of Ace2N-mNeon throughout zebrafish development from the gastrula stage, with membrane localization, and persisted across various cell types during early stages up to 5 dpf with the Tol2 transposon system.

### 2.2 Cell-Specific Constructs Enable Differential Ace2N-mNeon Expression in Developing Danio rerio

In our initial experiments, we observed ubiquitous expression of Ace2N-mNeon throughout zebrafish development. However, this broad expression made it difficult to distinguish between different cell types without a specific cell marker or detailed morphological inspection. To address this, we established a zebrafish Ace2N-mNeon developmental expression library, enabling the precise investigation of bioelectrical activity within distinct cell types throughout development (Table 1). Given the crucial roles of neurons and myocytes during development, our research mainly focused on these cell types. Neurons are essential for propagating bioelectric signals that orchestrate tissue and organogenesis^3^, while myocytes are integral to motility, structural support, and cardiac functionality^53^.

**Table 1.**

| Promoter | Target cell types |
| --- | --- |
| 503-bp promoter of zebrafish *unc-45b* | skeletal and cardiac muscle cells |
| α-Smooth Muscle Actin (*acta2*) | smooth muscle cells and pericytes |
| cardiac myosin light chain 2 (*cmlc2*) | cardiomyocytes |
| ELAV-like protein 3 (*elavl3*) | neurons |
| Neurogenic differentiation 1 (*neuroD1*) | neurons |

Zebrafish embryogenesis is marked by rapid development, with somite formation beginning around 16 hpf and the basic body plan and early organ systems forming by 19.5 hpf ^54^. To investigate bioelectricity changes during this crucial period, we analyzed promoter-driven expressions starting at the 21-somite stage (19.5 hpf), a key point for early organogenesis^54,55^. As expected, our constructs showed cell-specific expression, though expression efficiency varied, likely due to cell differences in DNA recombination across cell types (Figs. 2, 3).

**Fig. 2.**
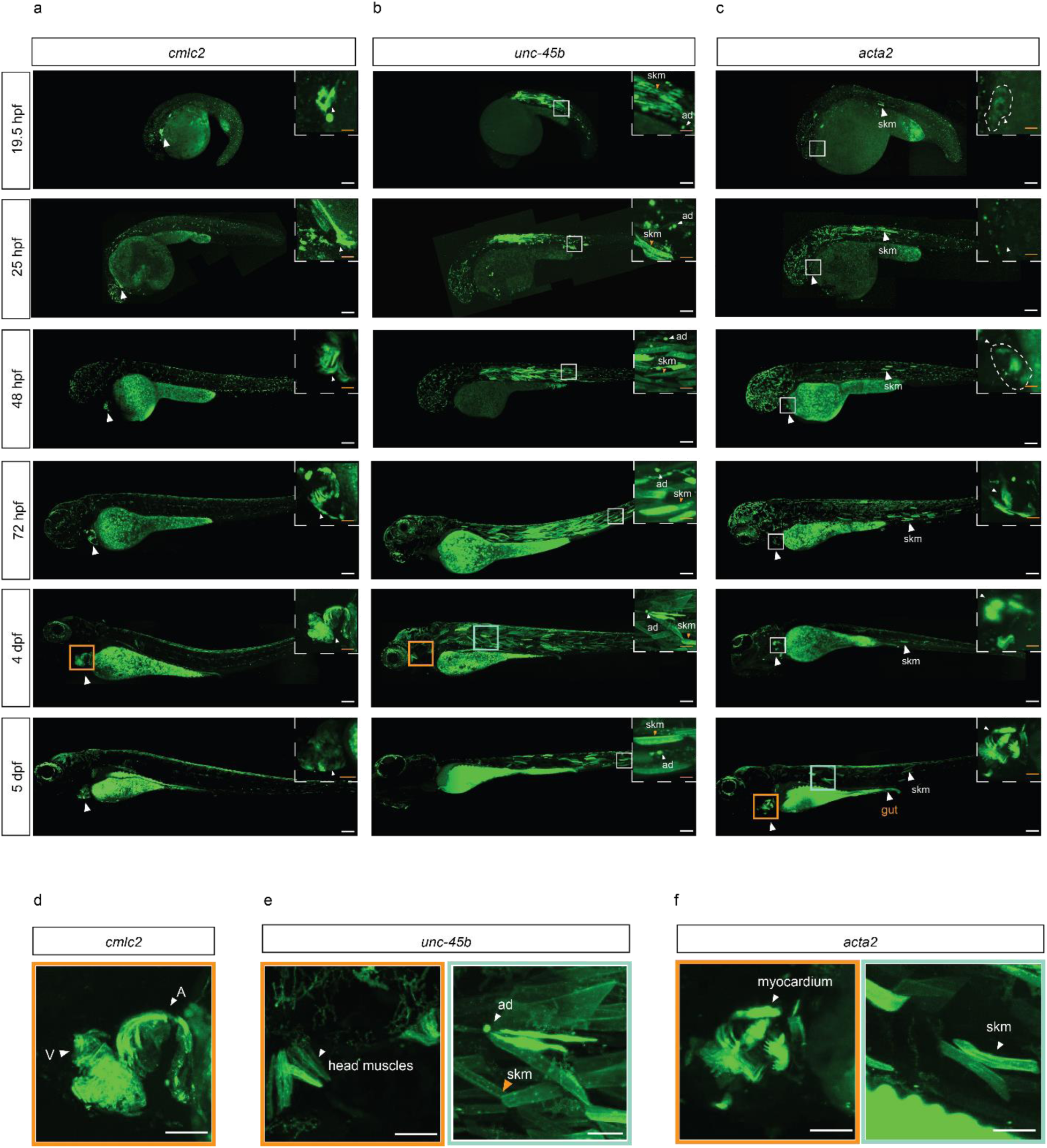
Ace2N-mNeon expression in muscle cells throughout developmental stages. (a-c) Confocal fluorescence images of Ace2N-mNeon expression in different muscle cells throughout development. Scale bar = 100 μm. Magnified views within dashed boxes have a scale bar of 25 μm. (a) Maximum projection of the entire zebrafish under cmlc2 promoter from 19.5 hpf to 5 dpf. Ace2N-mNeon expressed in cardiomyocytes from the 21-somite stage (19.5 hpf), with increasing intensity over time. (b) Maximum projection of the entire zebrafish under unc-45b promoter from 19.5 hpf to 5 dpf. Ace2N- mNeon displayed strong expression in skm (dashed boxes with orange arrow) and ad (dashed boxes with white arrow). Fluorescence is also evident in head muscles by 4 dpf (orange box). (c) Maximum projection of the entire zebrafish under acta2 promoter from 19.5 hpf to 5 dpf. Ace2N-mNeon expression is observed in a few skm at 19.5 hpf. From 48 hpf, expression becomes more visible in the myocardium, and by 5 dpf, strong expression is observed in both the myocardium (orange box) and the visceral smooth muscle of the gut (orange arrow). (d-f) Maximum projection of zoom-in image at later stages under the promoters. Scale bar = 50 μm. (d) Single z-plane confocal image of Ace2N-mNeon expression under the cmlc2 promoter. Ace2N-mNeon can be observed in cardiomyocytes, with clear fluorescence in the atrial (A) and ventricle (V). (e) Single z-plane confocal images of Ace2N-mNeon expression at 4 dpf under the unc-45b promoter. Ace2N-mNeon can be observed in the head muscles (orange box), skm (green box with orange arrow), and ad (green box with white arrow). (f) Single z-plane confocal images of Ace2N-mNeon expression at 5 dpf under the acta2 promoter. Ace2N-mNeon can be observed in the myocardium (orange box) and skm (green box). hpf: hours post-fertilization; dpf: days post-fertilization; skm: skeletal muscle cells; ad: adaxial cells.

**Fig. 3.**
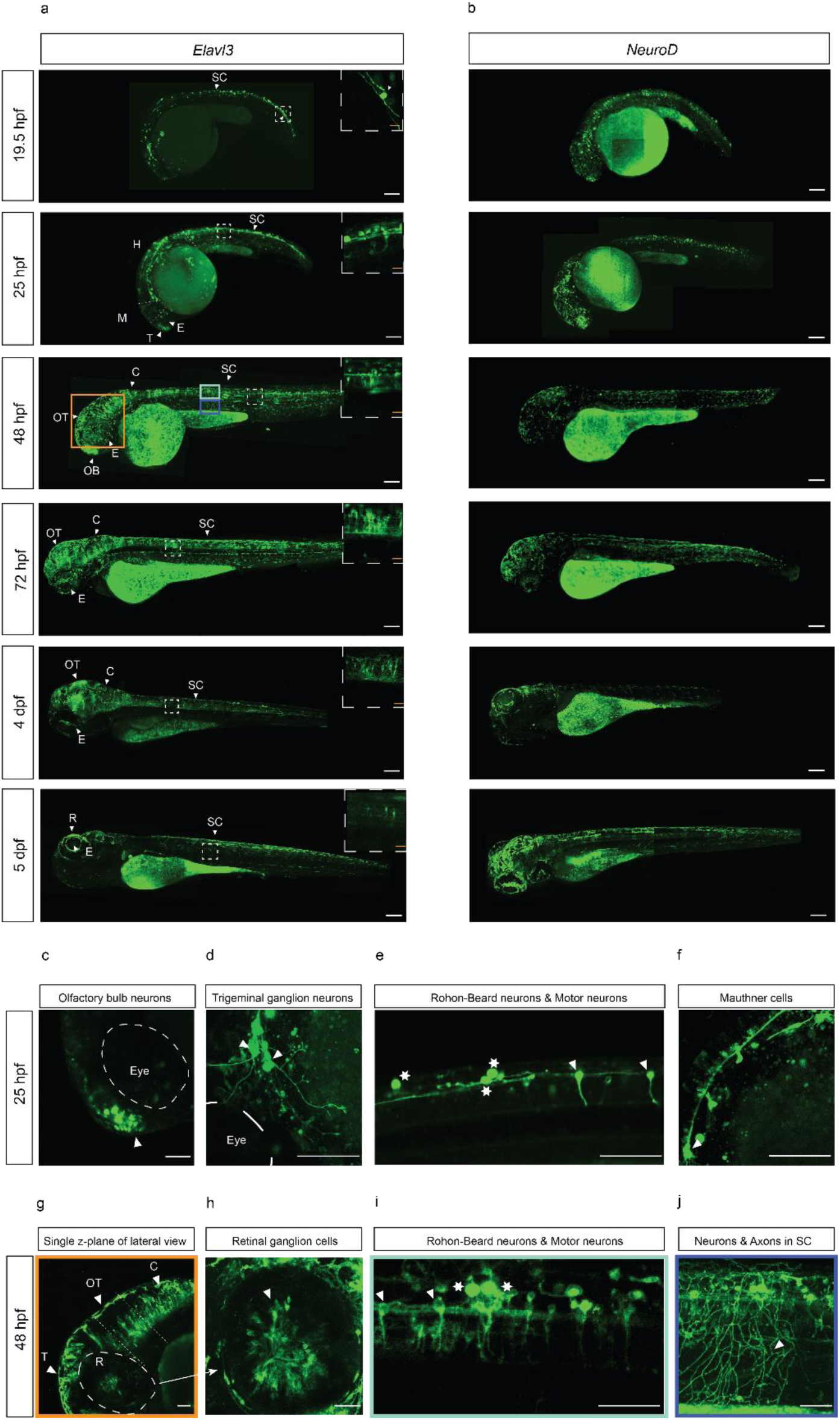
Ace2N-mNeon expression in neurons throughout developmental stages. (a-b) Confocal fluorescence images of Ace2N-mNeon expression in neurons throughout development. Scale bar = 100 μm. (a) Maximum projection of the entire zebrafish under elavl3 promoter from 19.5 hpf to 5 dpf. Prominent expression in the spinal cord (SC) can be observed at 19.5 hpf, with expression appearing in the telencephalon (T) and hindbrain (H) by 25 hpf, and increased expression in the olfactory bulb (OB), retina (R), optic tectum (OT), and cerebellum (C) from 2 dpf to 4 dpf. Scale bar = 100 μm. Single z-plane confocal images from a specific brain region are displayed in the white dashed-line box at the top right corner. Scale bar = 20 μm. (b) Maximum projection of the entire zebrafish under neuroD promoter from 19.5 hpf to 5 dpf. Expression levels are lower compared to the elavl3 promoter. (c-f) Single z-plane confocal images of Ace2N-mNeon expression in specific neuronal populations at 25 hpf. Neurons are identified by location and morphology. Scale bar = 50 μm. (c) Olfactory bulb neurons in the telencephalon (white arrows). (d) Trigeminal ganglion neurons (white arrows). (e) Rohon-Beard mechanosensory neurons (white stars) and motor neurons (white arrows). (f) Mauthner cells (white arrows). (g) Single z- plane confocal images of Ace2N-mNeon expression in the retina (R), optic tectum (OT), and cerebellum (C) at 48 hpf (area indicated in (a) by the orange box). Scale bar = 50 μm. (h) Single z-plane confocal images of Ace2N-mNeon expression in retinal ganglion cells (RGCs) at 48 hpf. The RGCs are distributed and dense in the retina. Scale bar = 50 μm. (i) Single z- plane confocal image of neurons and axons in the spinal cord at 48 hpf. A denser expression pattern can be observed in the spinal cord. The area indicated in (a) by the green box, Scale bar = 50 μm. (j) Single z-plane confocal image of Ace2N-mNeon expression in spinal cord axons at 48 hpf. The secondary motor neuron axons project to the ventral trunk muscles, illustrating their functional connectivity. Scale bar = 50 μm. **hpf**: hours post-fertilization; **dpf**: days post-fertilization.

#### 2.2.1 Ace2N-mNeon expression in muscle cells

Zebrafish musculature comprises smooth, skeletal, and cardiac muscle cells, each with distinct gene expression profiles and localizations^53^. The promoters used in this study allow the differentiation and independent analysis of these cell types.

Cardiac Muscle Cells: cardiac myosin light chain 2 (*cmlc2*) is a key contractile protein in cardiac and skeletal muscles^56^. For this study, we employed the zebrafish *cmlc2* promoter, which is transformed and allows specific expression in cardiomyocytes^57^. The expression begins at 19.5 hpf and continues throughout development (Fig. 2a). By 4 dpf, distinct structures of the atrial (A) and ventricle (V) are observable (Fig. 2d), enabling visual recording of the heartbeat.

Skeletal Muscle Cells: *unc-45b* is a myosin chaperone which is crucial for the correct assembly of the contractile apparatus in developing muscles^58–60^. In zebrafish, it is specifically expressed in striated muscle, including both cardiac and skeletal muscles, where it can help myosin fold during myofiber formation ^57–61^. At 19.5 hpf, the *unc-45b* promoter labels skeletal muscle cells (skm) and adaxial cells (ad), the progenitors of slow-twitch skeletal muscle cells (Fig. 2b, 19.5 hpf). These cells can be identified by their morphology (Fig. 2b, e). By 3 dpf, fluorescence was evident in head muscles (Fig. 2e).

Smooth Muscle Cells: α-Smooth muscle actin (*acta2)* is one of the earliest markers for mural cell development in vertebrates, expressed in smooth muscle cells (SMCs) and pericytes^62^. In zebrafish, *acta2*-EGFP expression begins slightly later in development, initially in the myocardium, followed by visceral smooth muscle and skeletal muscle, and eventually in vascular smooth muscle^62^ At 19.5 hpf, *acta2* expression is detectable in the myocardium, though not very clear, together with a few skeletal muscle cells (Fig. 2c, 19.5 hpf). More expression in skeletal muscle cells could be observed from 25 hpf onwards (Fig. 2c, 25 hpf). By 48 hpf, *acta2* expression becomes more visible in the myocardium (Fig. 2c, 48 hpf). By 5 dpf, strong expression was observed in the myocardium and visceral smooth muscle of the gut (Fig. 2c, f, 5dpf).

#### 2.2.2 Ace2N-mNeon expression in neurons

During early somite formation, *elavl3*, an mRNA-binding protein expressed in nearly all postmitotic or newly differentiated neurons^63^, is detectable in the neural tube and other brain regions of zebrafish embryos, while lower expression is observed in other brain regions (Fig. 3a). At 1 dpf, our results show that Ace2N-mNeon expression is prominently enriched in neurons of the spinal cord (SC) and other brain regions like telencephalon (T) (Fig. 3a, 19.5 hpf, 25 hpf). Notably, several neuronal populations, including olfactory bulb neurons^64,65^, trigeminal ganglion neurons^66^, mauthner cells^67^, Rohon-Beard mechanosensory neurons^66,68^, and motor neurons^69^, can be clearly identified at this stage, most of which exhibit soma or small axons (Fig. 3c-f).

As development progresses to 2 dpf, Ace2N-mNeon expression expands throughout the Central Nervous System (CNS). In addition to the brain regions observed at 1 dpf, expression becomes visible in the retina (R), optic tectum (OT), and cerebellum (C) (Fig. 3a, g). At this stage, retinal ganglion cells (RGCs) can be identified, with axons extending centrally (Fig. 3h), and secondary motor neurons projecting axons into the ventral trunk muscles also become distinguishable (Fig. 3i, j). From 2 dpf to 4 dpf, as neurons mature and express Ace2N-mNeon, clusters of neurons begin to appear in different regions, such as the optic tectum, cerebellum, and spinal cord, marking the early stages of functional circuit formation (Fig. 3a). The increased labeling density beyond 48 hpf reduces the visibility of individual neurons in maximum intensity projections, though they can be distinguished in single z-planes from 3D confocal scans (Fig. 3a, white dashed line box), Ace2N-mNeon expression persists in the CNS through at least 5 dpf.

Additionally, we compared Ace2N-mNeon expression under the *elavl3* promoter with expression under the *neuroD* promoter, another neuron-specific driver (Fig. 3a, b)^70^. Our findings indicate that *elavl3*-driven expression is denser than *neuroD*-driven expression (Fig. 3a, b), likely reflecting the physiological differences in their expression profiles. While *elavl3* is broadly expressed in almost all newly differentiated neurons in zebrafish embryos^71^, *neuroD* expression can have a lower or absent expression in certain neuron types^72,73^. Altogether, these results indicate that Ace2N-mNeon constructs allow cell-specific expression with membrane localization throughout zebrafish’s early development.

### 2.3 Ace2N-mNeon voltage imaging reveals neurogenesis-specific electrical activity

Bioelectricity comprises the electrical phenomena emerging from ionic currents that lead to voltage changes in the cellular membrane. In excitable cells, such as neurons or muscle cells, these ionic currents can create fast and large voltage fluctuations, also known as APs^5,74^, which can be studied using electrophysiology or imaging techniques ^22,75^. In vivo voltage imaging of multiple neurons is a powerful approach for characterizing neural dynamics, offering diverse information and insights into possible circuit mechanisms ^22,30,41,76,77^. To investigate whether Ace2N-mNeon can detect spontaneous activity in various neural populations during early development, we co-injected the pTol2pA2_elavl3_Ace2N- mNeon construct with Tol2 transposases mRNA into one-cell-stage embryos. Positive embryos were selected for high-speed imaging at different zebrafish brain regions from 24 to 28 hpf.

As shown in Figure 3, we identified several labeled neurons in each positive embryo. For further voltage imaging, we focused on specific neuron types based on their morphology: Rohon-Beard (RB) mechanosensory neurons, motor neurons, and terminal nerve cells in the olfactory bulb. Fig. 4b shows a dorsal view of the spinal cord, highlighting the labeled RB somas and their longitudinal axons, as well as spinal cord motor neurons with their distinct axons extending into the ventral myotome^69^, parallel to the RB neuron axons. Olfactory bulb nerve cells are located near the axon bundles of the medial olfactory tract in the rostral portion of the bulb, close to the zebrafish eyes (Fig. 4c)^65^. We recorded neural signals from these neuron types as early as 25 hpf (Fig. 4d-f). Furthermore, Fig. 4g and 4f show that Ace2N-mNeon enables populational neural recordings of different neuron types within a single field of view.

**Fig 4.**
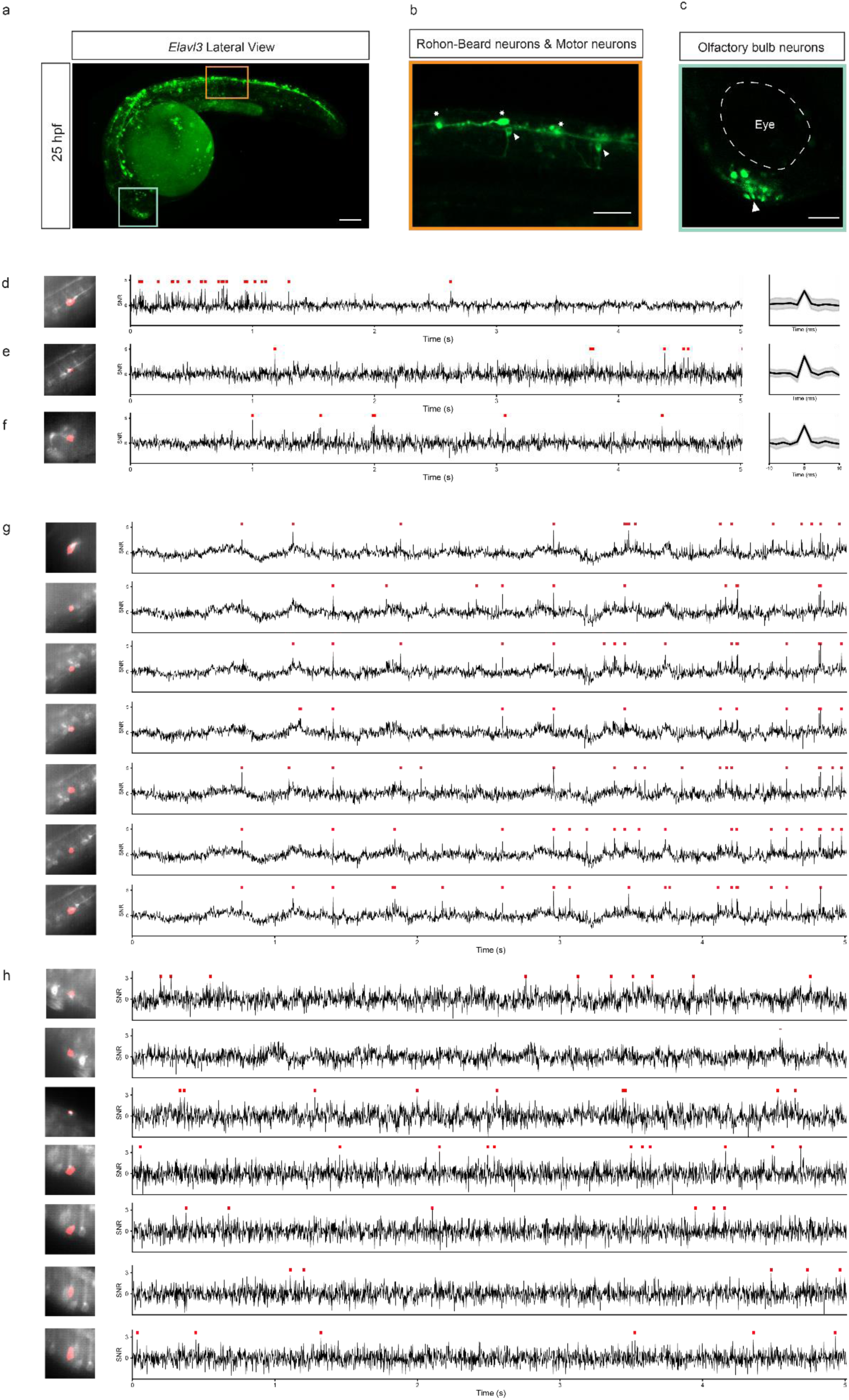
Voltage imaging with Ace2N-mNeon enables the detection of electrical activity in various neuron types across developmental stages. (a-c) Confocal images of zebrafish at 25 hpf. Scale bar = 100 μm. (a) Maximum projection of the entire zebrafish. Orange and green squares indicate regions for a magnified view, as shown in (b) and (c). (b) The single z-plane of the confocal image showing motor and Rohon-Beard (RB) neurons is (white arrows). (c) Single z-plane of the confocal image shows an olfactory nerve cell cluster. (d-g) Electrical activity recordings of neurons. Scale bar = 100 μm. (d) Motor neuron. (e) RB neuron. (f) Olfactory terminal nerve cell (g). The left panels show the average image of the recording with the spatial footprint. The center panels show the example traces. The right panels show the spike waveform averages with standard deviation based on the detected spikes (indicated in the traces). (h-i) Electrical activity of multiple motor neurons recorded in the same field of view at 24 hpf (h) and 3 dpf (i). The left panels show the average image of the recording with spatial footprints, manually annotated in orange. For all traces, red ticks indicate detected spikes.

Studying membrane voltage changes provides insights into both neural activity and neurodevelopment. During zebrafish development, neural progenitor cells undergo primary neurogenesis to form a neuronal scaffold, followed by secondary neurogenesis where axons and dendrites extend and first synapses form^78–80^. To investigate how neural activity develops, we performed voltage imaging in motor cortex populations at 3 dpf. As neurons mature and express the indicator, labeling becomes denser (Fig. 3a, i), and it is difficult to image individual cell imaging at 3 dpf due to the brighter background. However, by focusing on sparsely labeled areas, we successfully identified and imaged individual cells. Notably, putative motor neuron activity at 25 hpf (Fig. 4g) was more synchronized than at 3 dpf (Fig. 4h). Additionally, we recorded activity in previously unidentified neuron types (Supplementary Fig. 3a-c). These findings indicate that our approach effectively studies neural activity throughout development.

### 2.4 Detection of voltage dynamics in the zebrafish heart in vivo

Zebrafish is an ideal model for investigating cardiac development and human cardiac diseases^81^, as 96% of human cardiomyopathy-related genes are retained and highly expressed in this model organism^82^. In our study, the expression library driven by the *cmlc2* promoter showed strong localization to the plasma membrane of cardiomyocytes from the 21-somite stage, persisting throughout development (Fig. 2a). Cardiac atria and ventricles were clearly distinguishable at 48hpf (Fig. 2a, 48 hpf). To track zebrafish cardiac AP changes during early development, we recorded fluorescence of the zebrafish heart on *cmlc2*: Ace2N-mNeon-positive zebrafish embryos at 25 hpf, 48 hpf, and 102 hpf (Fig. 5c-e). During early development, zebrafish can remain healthy even with inhibited heart contractions, as oxygen diffuses passively through tissues^81^.

**Fig. 5.**
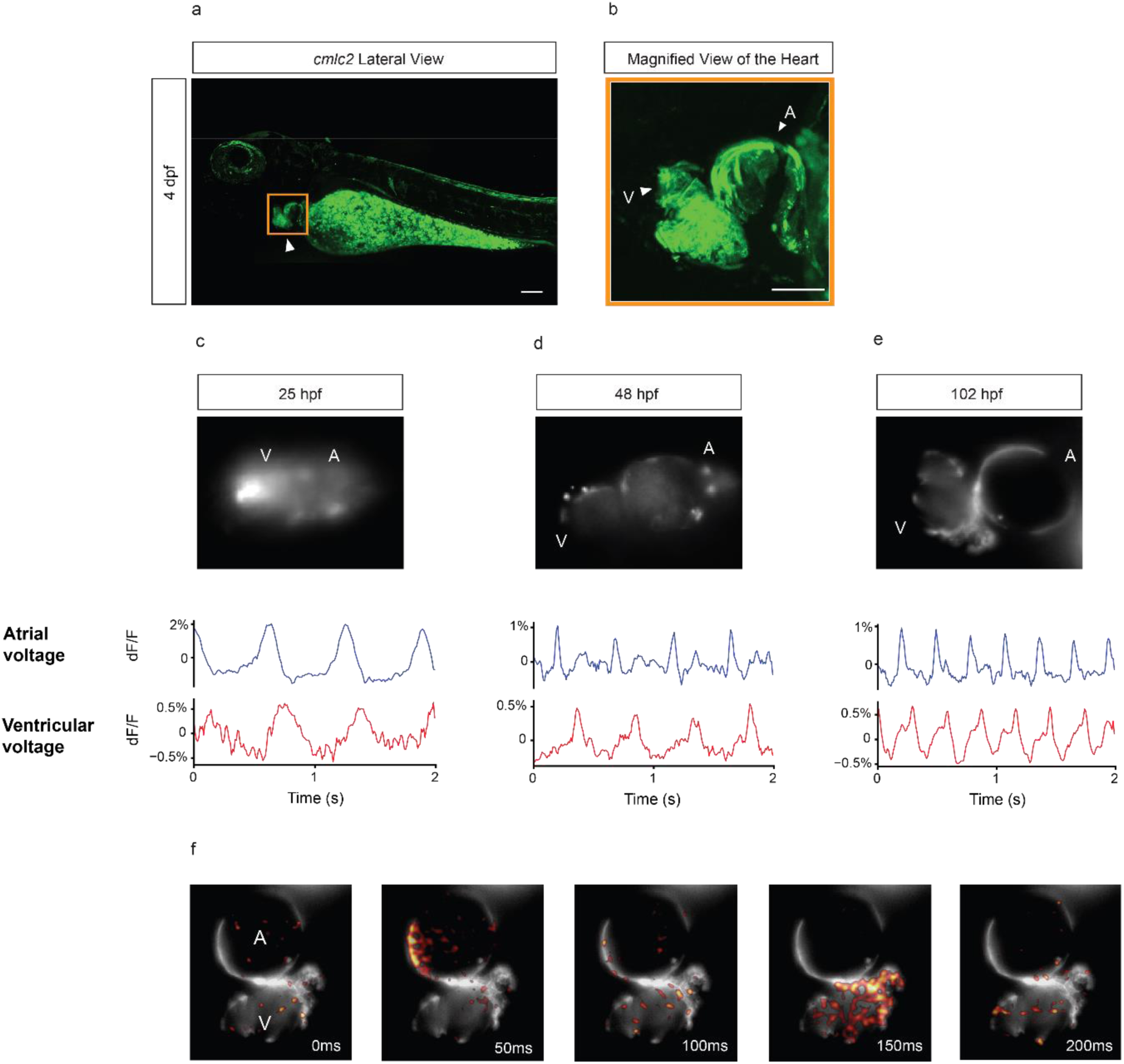
Voltage transients of the atrial and ventricle in an embryonic zebrafish heart during development. (a) Confocal image of the zebrafish heart at 4 dpf under the cmlc2 promoter. Scale bar = 100 μm. (b) Magnified view of the zebrafish heart expressing Ace2N-mNeon at 4 dpf. Scale bar = 50 μm. (c-e) Voltage transients in the atria and ventricles of the zebrafish embryonic heart during development. Top: Fluorescence images of the positive zebrafish embryonic heart for Ace2N- mNeon. At 25 hpf, the heart is tubular, with no clear separation between the ventricle and atrium. By 48 hpf, the two chambers are distinctly visible. Bottom: Voltage imaging wavelengths of Ace2N-mNeon in the atrial (A) and ventricle (V) at (c) 25 hpf, (d) 48 hpf, and (e) 102 hpf. At 25 hpf, lower Ace2N-mNeon expression leads to higher noise compared to other time points. At all-time points, blue light intensity was minimized to reduce the photoinactivation of para-aminoblebbistatin. (f) Representation of zebrafish heart voltage dynamics at 102 hpf. Colored representation of positive voltage changes from the baseline, following atrial-to-ventricle voltage dynamics. **hpf**: hours post-fertilization; **dpf**: days post-fertilization.

To minimize motion artifacts from heart contractions, embryos were treated with 100 µM para-amino blebbistatin 4 hours before imaging to inhibit the heartbeats^83^. We found that para-aminoblebbistatin is sensitive to blue light, after each recorded zebrafish, we increased the light intensity to restore heart contractions, confirming embryo viability. Upon exposure to blue light, the heart resumed strong contractions at all development stages, indicating healthy conditions (Supplementary Movie. 1-3).

We mapped the spatial and temporal progression of voltage patterns at 25, 48, and 102 hpf, using separate fish for each time point (Fig. 5c-ef). At 25 hpf, although the zebrafish heart appeared as a narrow tube with no clear division between atrial and ventricular chambers (Fig. 5c top), distinct voltage changes characteristic of atrial and ventricular AP waveforms was still detectable in different regions of the heart, with AP propagation in an atrial-to-ventricular direction (Fig. 5c bottom). It is important to note that the lower Ace2N-mNeon expression at this stage resulted in higher noise in the waveforms compared to later time points.

Starting at 48 hpf, the compartmentalization of the atria and ventricles became clear, with high fluorescence expression leading to much clearer waveforms and distinct voltage changes (Fig. 5d-e). AP propagation from the atria to the ventricle was still observed, and these voltage changes became faster as development progressed, reflecting the increased zebrafish heart rate (Fig. 5d-e). By 102 hpf, APs in the atria and ventricle occurred as two clearly resolved beats, and Fig. 5f shows the trajectory of voltage propagation from the atrium to the ventricle. Notably, the electrical APs we recorded optically in vivo were consistent with previous findings from patch-clamp measurements on explanted hearts^84,85^.

## 3. Discussion & Conclusion

GEVIs have revolutionized the study of bioelectric signals, enabling both fast and slow voltage changes to be monitored through different imaging modalities. In this work, we selected Ace2N-mNeon for its high brightness, rapid kinetics, and measurable fluorescence lifetime^48^, which allows for detecting not only voltage changes but also absolute membrane voltage values^34^. To our knowledge, this work presents the first demonstration of Ace2N- mNeon expression across multiple cell types in zebrafish^86^. To achieve this, we developed a library of early expression plasmids for zebrafish, enabling expression starting from the 512- cell stage. This library supports both ubiquitous and cell type-specific expression in neurons, cardiomyocytes, mural cells, and skeletal muscle cells, enabling the study of bioelectric signals across different cell types and developmental stages (Supplementary Table S2). Although ubiquitous expression may introduce fluorescence crosstalk and be less effective for single-cell studies, it remains valuable for examining cellular and tissue-level bioelectric signaling, especially when cell-specific markers are unavailable.

A key application of this library is tracking spiking activity in the early-developing brain via voltage imaging. Zebrafish neurogenesis can be divided into primary neurogenesis (up to 2dpf), characterized by the formation of a transient neuronal scaffold, and secondary neurogenesis, which involves further differentiation of neurons and enhanced dendritic and synaptic growth^79,80,87^. While previous studies have correlated GEVI fluorescence with fast-spiking activity, few have explored such activity during early development^28^. Voltage imaging using GEVIs allowed us to explore the relationship between neural activity and neurogenesis, revealing distinct neuronal activity at different developmental stages. Developing neural networks often exhibit synchronous oscillatory activity, especially during periods of intense axonal growth and synaptogenesis^80,87^. Our results validate the use of this library to study dynamic changes during neurogenesis, revealing synchronous oscillatory activity in spinal neurons as early as 20 hpf^88^. This synchronicity, associated with spontaneous contractions, was not observed at later stages, aligning with existing literature suggesting that synchronized activity is specific to early neurogenesis^88^. These findings underscore the potential of voltage imaging to capture neurogenesis-dependent electrical phenomena with high spatiotemporal resolution.

Despite its advantages, voltage imaging has several limitations. The short duration of spikes (< 4 ms) poses a challenge for their detection that depends on the indicator’s kinetics, expression levels, and the recording setup^89^. Additionally, fluorescence signals can be contaminated due to fluorescence fluctuations of neighboring cells^77^, especially in one-photon imaging, where out-of-focus fluorescence can decrease signal-to-noise ratios (SNR) of the collected traces. This issue becomes even more pronounced in densely labeled zebrafish embryos, due to their transparency. The broad expression of the promoters used (Elavl3 and NeuroD) in a wide range of neurons may explain why cell activity is more easily identified at 24 hpf but less so at 3 dpf^70,90^. To improve SNR at later developmental stages, researchers can label specific neural populations with cell-specific promoters in future work. Additionally, incorporating sub-cellular targeting motifs, such as the commonly used Kv2.1 for somatic targeting, could significantly reduce background fluorescence from dendrites and axons^91^.

While voltage imaging developments have predominantly focused on recording fast neural signals, GEVIs are also suitable for detecting slower APs, such as those in cardiomyocytes. Previous studies have demonstrated this in iPSC-derived cardiomyocytes^92^, excised hearts^93^, and in vivo zebrafish^29^. Here, we show that atrial and ventricular voltage activity can be distinguished even earlier before a clear anatomical division of heart chambers is formed. Although no definitive conclusions can be made, this data, together with the role of bioelectricity in patterning during tissue formation, might suggest a potential relationship between electrical activity and heart morphological development.

Beyond fast-spiking signals, the role of slower RP changes in development remains underexplored. Non-excitable cells use their ionic properties as endogenous guidance cues that regulate individual cell behavior^9^, organ shape^14,15,18^, and organismal development^19^. Understanding these signals is crucial for elucidating growth and morphogenesis regulation and developing targeted treatments^2,4,6,7^. Current research focuses on changing bioelectricity by modulating membrane potential (Vm) through ion channel manipulation, which can induce specific developmental changes^2,14,94^, reverse teratogen-induced brain defects^95^, and neurogenesis-related genetic mutations^19^. These pathways offer promising targets for biomedical strategies addressing birth defects, regeneration, and cancer suppression. Although some research has successfully recorded RP changes in cells^34,35,86,96^, current tools are not yet optimized for capturing slow, long-term electrical signals in live organisms. Our Ace2N-mNeon library demonstrates, for the first time, the potential of using GEVIs for in vivo recording of RP changes during development. These findings pave the way for future studies aimed at elucidating the bioelectric basis of growth and morphogenesis. Continued advancements in optical methods for precise, long-term measurement of RP changes, combined with our expression library, will further expand research opportunities across diverse cell types.

In summary, our research demonstrates the effectiveness of Ace2N-mNeon in studying bioelectricity during early biological development. This study highlights Ace2N-mNeon’s ability to monitor both rapid and long-term bioelectric changes in a non-invasive, real-time manner, positioning it as a versatile tool for broader applications in developmental and neurological research. Future research should refine this technique for other model organisms and explore its application in disease models to investigate the bioelectrical basis of developmental disorders. Expanding the toolkit for bioelectric studies, our work paves the way for deeper understanding and new discoveries in developmental biology and neurobiology.

## 4. Materials and Methods

### Zebrafish husbandry

All experiments were conducted using zebrafish (Danio rerio) of the *Casper* genotype^39^. Adult zebrafish were housed in recirculating systems at the Delft University of Technology fish facility, maintained under a 14-hour light/10-hour dark cycle, with a pH of 7.5 and a temperature of 28°C.

### Plasmid Construction

Ace2N-mNeon was amplified from YA1611: mNeon-Ace (a gift from Adam Cohen) and fused with various promoters using the Tol2 transposon system. This fusion was achieved via overlap-extension PCR using Phusion high fidelity master mix (NEB) DNA polymerase. The backbone containing the distinct promoters was amplified using KOD Xtreme Hot Start DNA Polymerase (Merck Sigma). The Ace2N-mNeon fragments and the promoter-containing backbone were then assembled using Gibson Assembly (Gibson Assembly Master Mix, NEB). The final assembly was transformed into NEB® 5-alpha Competent cells. The primers used for cloning are listed in the Supporting Information (SI, Table S1), and all the plasmids we constructed are listed in the Supporting Information (SI, Table S2).

### RNA Synthesis

Ace2N-mNeon was first amplified from YA1611: mNeon-Ace and fused with SP6 promoter from pCS2FA-CO-Tol2-TPase (Addgene: #133032). Tol2 transposase mRNA was transcribed directly from pCS2FA-CO-Tol2-TPase. RNA synthesis was then performed using the mMESSAGE mMACHINE® Kit (Thermo Fisher Scientific). In vitro transcription was conducted according to the manufacturer’s protocol to ensure high-quality RNA production. The synthesized RNA was then purified and quantified before use in subsequent microinjections.

### Microinjection

Fertilized zebrafish eggs were collected at the 1-cell stage immediately after spawning. Microinjections of Ace2N-mNeon mRNA (100 ng/µL) with 0.1% phenol red (Sigma-Aldrich, P0290) were performed at the 1-4 cell stage, injecting approximately 1 nL of mRNA solution into the cytoplasm. For DNA injections, 1-cell-stage embryos were co-injected with 1 nL of a solution containing 50 ng/µL plasmid DNA, 100 ng/µL Tol2 transposase mRNA, and 0.1% phenol red into the yolk. After injection, embryos were incubated in egg water containing instant ocean salts (60 μg/ml), and 0.0002% methylene blue, and unfertilized or dead eggs were removed at 6- and 24-hours post-injection.

### Confocal Imaging Sample preparation

For confocal imaging, embryos from 2.75 hpf to 12 hpf were imaged directly with one drop of egg water. Embryos from 19.5 hpf to 5 dpf were manually dechorionated (before 2 dpf) and mounted in 1.5% agarose (Sigma-Aldrich, A4018) with 0.03% tricaine (Sigma-Aldrich, A5040). Before and after imaging, embryos were visually inspected for a heartbeat, overall health, and the absence of developmental abnormalities.

### Confocal Imaging Setup

Confocal imaging was conducted on a Nikon Ti C2si inverted confocal microscope using a 485 nm excitation wavelength with an illumination power of 3.35 mW. The system was equipped with a 405/488/561/640 dichroic mirror (Nikon, MHE46410), only the 488 nm dichroic mirror was used in this study. A 525/50 emission filter (Nikon, MHE46710) was employed. Fluorescence images were captured using a 20x objective (Nikon CFI Plan Fluor 20XC MI, NA 0.75). Data acquisition was performed with Nikon’s NIS-Elements software, and subsequent image analysis was carried out using ImageJ.

### Voltage Imaging of Neurons

Embryos aged 24-28 hpf and 3 dpf were paralyzed with 1 mg/mL α-Bungarotoxin (Sigma-Aldrich, 203980) for 15 minutes to block nicotinic acetylcholine receptors and prevent motion artifacts during imaging. They were then mounted in 1.5% low-melting agarose (Sigma-Aldrich, A4018) for single-photon imaging, performed on an A1RMP Nikon microscope with a 25x objective (Nikon CFI75 Apo, 25XC W, NA 1.1). Recordings were made using a white LED light source (SOLA Light Engine, Lumencor) at 20-50 mW/mm², filtered with a 504/12 excitation filter (Semrock FF01-504/12-25), 518 high-pass dichroic mirror (Semrock FF518-Di01-25x36), and 532/18 emission filter (Semrock FF01-532/18-25). Images were acquired at 500 frames per second (fps) using a Kinetix sCMOS camera (Teledyne Photometrics) controlled by μManager, with 2x2 or 4x4 spatial binning, for 20-30 seconds.

### Neuronal voltage imaging analysis

Voltage imaging recordings first corrected the motion artifacts using a modified version of the rigid motion correction algorithm from NoRMCorr within the CaImAn infrastructure^97–99^. Next, motion-corrected videos were denoised using the SUPPORT pipeline^99^, with a model trained for 100 iterations on zebrafish tail recording of 5000 frames (10 seconds). Following denoising, regions of interest (ROIs) corresponding to individual cells were manually annotated and processed using the Volpy pipeline^100^. Volpy incorporates parameters for refining spatial footprints and extracting signals, with most settings maintained at default values except for the high-pass filter set to 1 Hz to optimize spike detection. Cells exhibiting well-defined spatial footprints post-Volpy processing were selected for subsequent analysis.

For data visualization in Figure 4, all traces were lastly subjected to a 1Hz high-pass filter for baseline correction, ensuring clarity in depicting neural activity dynamics. All traces are shown as SNR, calculated by the peak of the spike divided by the baseline standard deviation.

### Voltage Imaging of Heart Dynamics

Embryos at 25 hpf, 48 hpf, and 102 hpf were reared in egg water. The 25 hpf and 48 hpf embryos were manually dechorionated and incubated with 100 μM para-aminoblebbistatin (MCE, HY-111474) for 4 hours to inhibit heart contractions. Following incubation, the embryos were mounted in 3% low-melting agarose (Sigma-Aldrich, A4018) for imaging. A drop of para-aminoblebbistatin was added 5 minutes before recording. For voltage imaging, single-photon imaging was performed on an A1RMP Nikon microscope with a 25x objective (Nikon CFI75 Apo, 25XC W, NA 1.1). Recordings were made using a white LED light source (SOLA Light Engine, Lumencor) at 0.6-1.5 mW/mm², filtered with a 504/12 excitation filter (Semrock FF01-504/12-25), 518 high-pass dichroic mirrors (Semrock FF518-Di01-25x36), and 532/18 emission filter (Semrock FF01-532/18-25). Images were acquired at 100 frames per second (fps) using a Kinetix sCMOS camera (Teledyne Photometrics) controlled by μManager, with 2x2 spatial binning, for 10 seconds each recording. For the representations of Figure 5, voltage imaging traces were filtered by a 1.5s Rolling Mean for photobleaching correction, and posterior 30 ms Rolling Mean for trace smoothing. To verify that zebrafish were still healthy, and restart heart contractions, 10 mW/mm² of the same light wavelength was shined upon the zebrafish heart for 10 seconds.

### Use of AI

ChatGPT and Grammarly were used to check phrasings of specific sentences in a first draft of this manuscript, before complete revision by authors. Any intellectual contributions, final decisions on content and structuring of the work, and writing of the final manuscript were exclusively made and done by credited authors.

## Acknowledgments

We gratefully acknowledge Huma Safar, Mariska Ouwehand, Aleksandra Placzek, Eleonora Muñoz-Ibarra, and Mike Lindhout for their technical support during the research that led to the writing of this paper. We thank Xin Meng and George Flamourakis for the voltage imaging data that was not included in this paper. We thank Elizabeth Carroll, Laura Maddalena, Byung Min Park, Marco Locarno, Marco Post, Bram Willems, and the Carroll lab in general for zebrafish husbandry during this project. DB acknowledges support from an NWO Start-up Grant (740.018.018) and ERC Starting Grant (850818 - MULTI-VIsion), as well as an NWO XS grant (OCENW.XS2.033) and by the Delft AI initiative (Ailab BIOlab) and the Convergence Health and Technology (Integrative Neuromedicine flagship).

## Author Contribution

**Conceptualization:** Daan Brinks, ZhenZhen Wu, Rui Oliveira Silva. **Plasmids preparation:** ZhenZhen Wu (lead), Srividya Ganapathy, Ruya Houssein, Fabiola Marques Trujillo, Jordan Gotti. **Experiment in expression library**: ZhenZhen Wu (lead), Ruya Houssein. **Expression library data analysis** & **Visualization**: ZhenZhen Wu (lead), Ruya Houssein. **Experiments in voltage imaging:** Rui Oliveira Silva (lead), ZhenZhen Wu. **Voltage imaging analysis & Visualization:** Rui Oliveira Silva (lead), ZhenZhen Wu. **Resources:** Daan Brinks (lead), Zhenyu Gao. **Funding, project administration, and supervision:** Daan Brinks. **Writing— original draft preparation:** ZhenZhen Wu, Rui Oliveira Silva, Daan Brinks. **Writing—review and editing:** Daan Brinks (lead), ZhenZhen Wu (lead), Rui Oliveira Silva (lead), Ruya Houssein, Fabiola Marques Trujillo, Jordan Gotti, Srividya Ganapathy, Zhenyu Gao. All authors have read and agreed to the published version of the manuscript.

## Ethics and inclusion statement

The research in this paper does not involve local researchers. All biological material was obtained according to the Nagoya protocol.

## Data availability statement

All data underlying manuscript figures have been uploaded to the 4TU Respository and are available at DOI: 10.4121/b9c28138-c971-455e-b4de-c8f29b1daa96

## Code availability statement

Voltage imaging analysis was performed with publicly available SUPPORT and VOLPY code

---

**Fig.S1.**
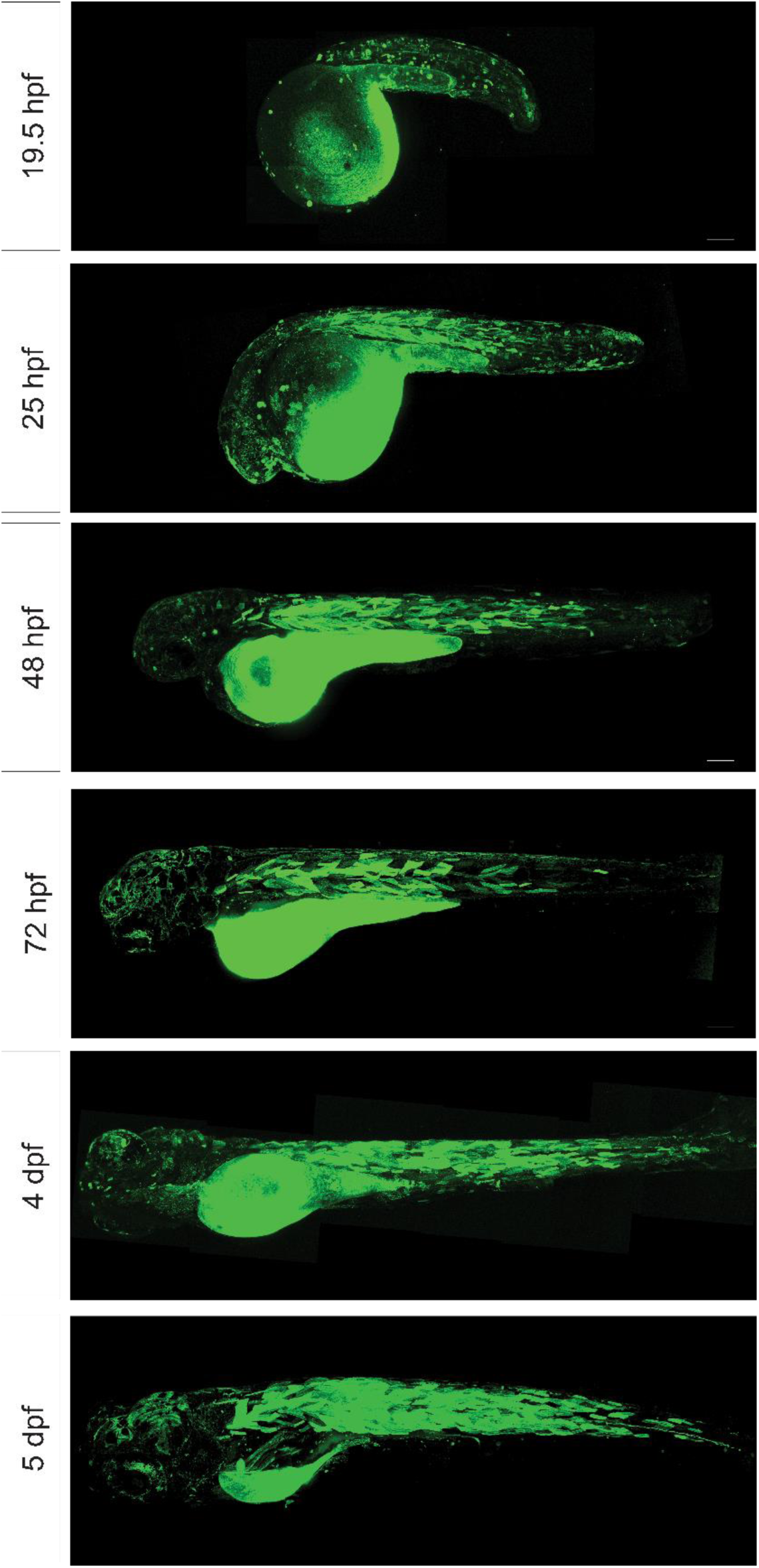
High-magnification images of Ace2N-mNeon expression driven by ubiquitin promoter.

**Fig. S2.**
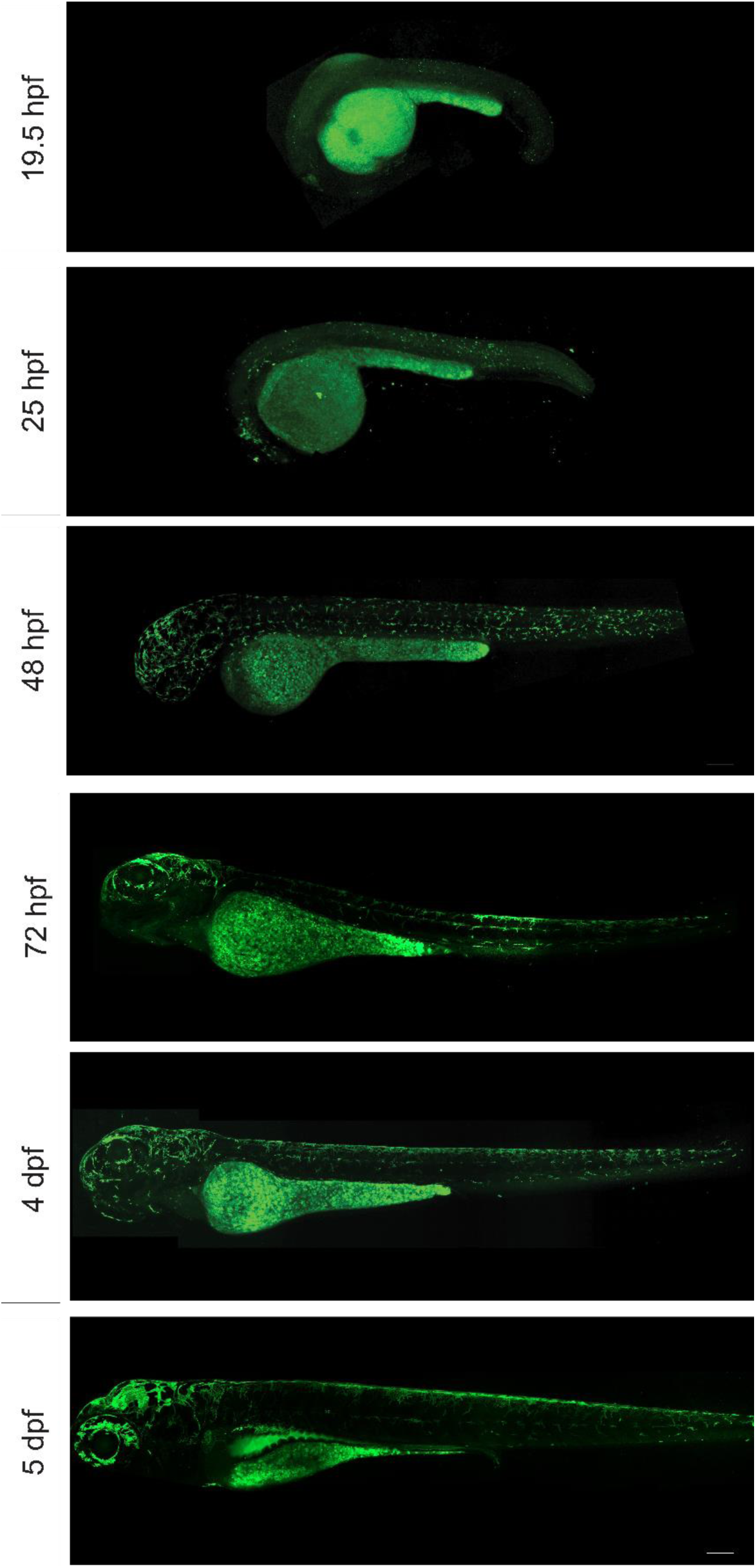
Autofluorescence in uninjected fish throughout development.

**Fig. S3.**
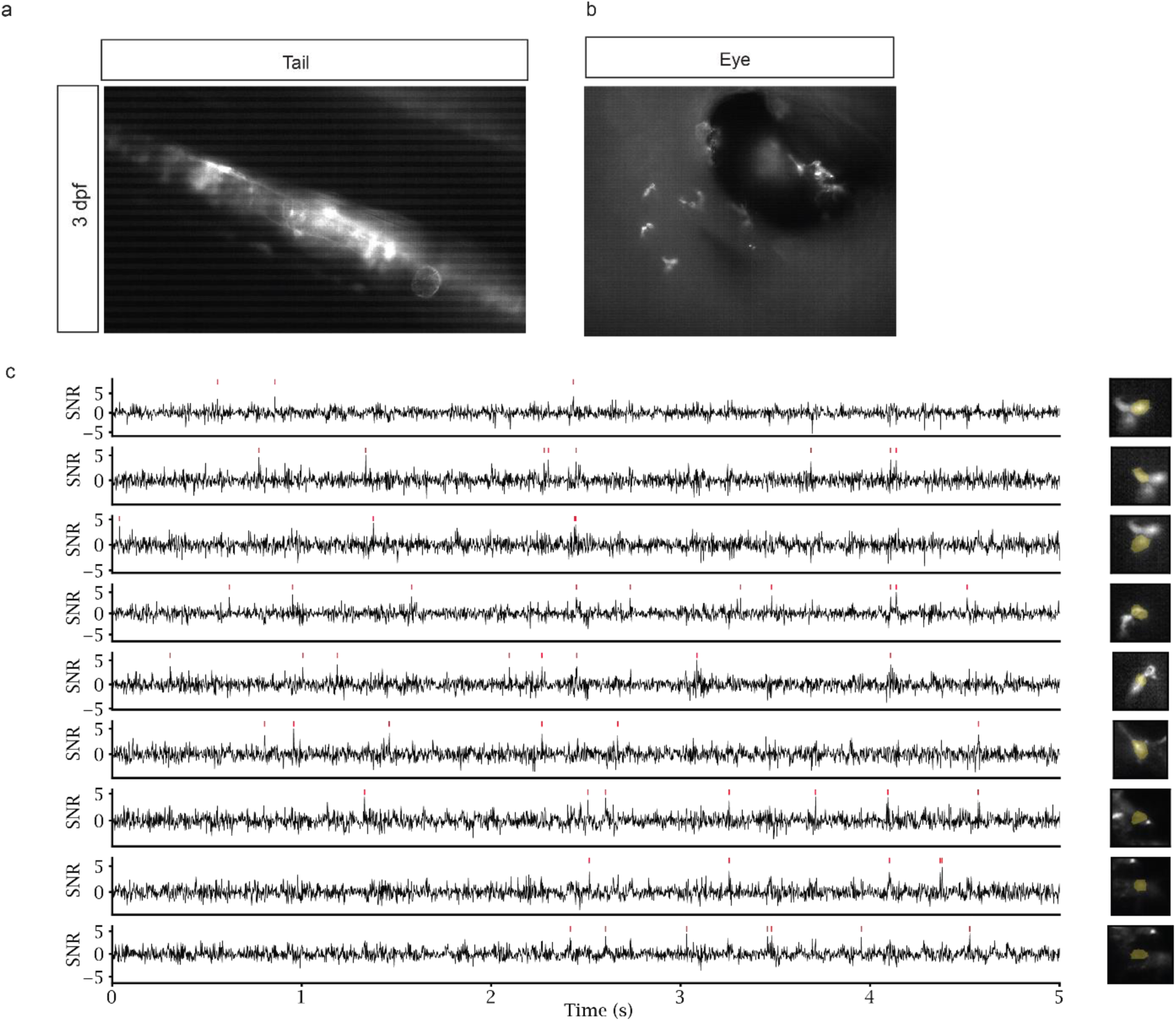
Voltage Imaging of Unrecognizable Neurons. (a-b) Representative images of fields of view from tail and head at 3dpf. (a) A single-photon image of the neurons in the tail shows a high background signal that makes them unsuitable for voltage recording. (b) A single-photon image of the neurons in the head, where the neurons near the eye are unrecognizable based on their morphology. (c) Representation of electrical activity of several neurons from b. For all traces, red ticks indicate detected spikes. Images on the right panels represent the average image of the recording, with a manually annotated spatial footprint in shaded orange.

**Table. S1.**
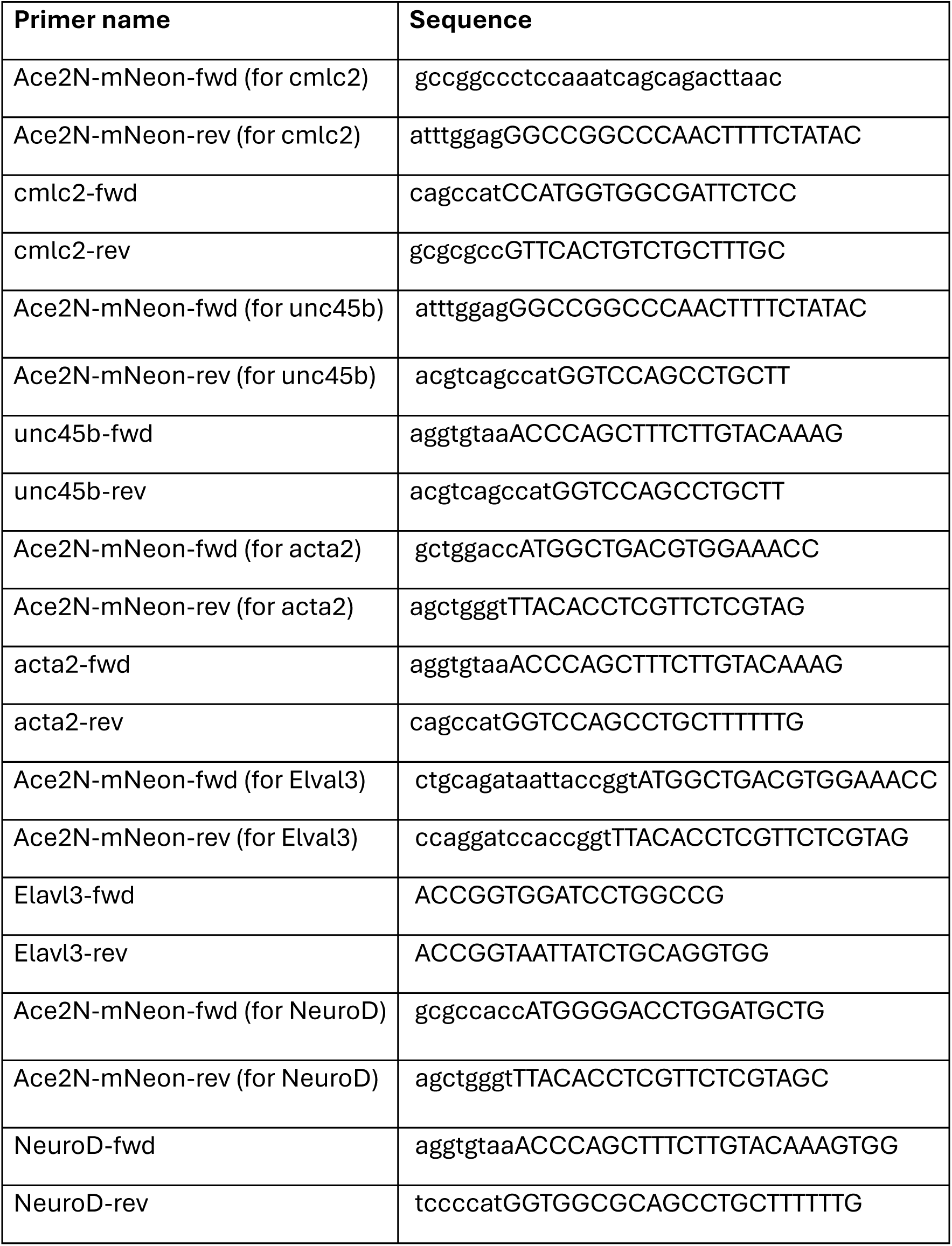
Primer sequences were used in this study. Primers used for Gibson assembly cloning of Ace2N-mNeon into the expression construct used in this study.

**Table. S2.** Plasmids were constructed in this study.

| **Plasmid name** |
| --- |
| pTol2pA2_ubi_Ace2N-mNeon |
| pTol2pA2_cmlc2_Ace2N-mNeon |
| pTol2pA2_punc503_Ace2N-mNeon |
| pTol2pA2_Acta2_Ace2N-mNeon |
| pTol2pA2_elavl3_Ace2N-mNeon |
| pTol2pA2_neurod_Ace2N-mNeon |

## Video S1-S3

**Video. S1. Restoration of Heart Contractions by Blue Light Exposure at 25 hpf. Video. S2. Restoration of Heart Contractions by Blue Light Exposure at 48 hpf. Video. S3. Restoration of Heart Contractions by Blue Light Exposure at 102 hpf.**

## References

1. Schofield, Z. et al. Bioelectrical understanding and engineering of cell biology. J R Soc Interface 17, 20200013 (2020).

2. Levin, M. Molecular bioelectricity in developmental biology: New tools and recent discoveries. BioEssays 34, 205–217 (2012).

3. Levin, M. Molecular bioelectricity: How endogenous voltage potentials control cell behavior and instruct pattern regulation in vivo. Mol Biol Cell 25, 3835– 3850 (2014).

4. George, L. F. & Bates, E. A. Mechanisms Underlying Influence of Bioelectricity in Development. Frontiers in Cell and Developmental Biology vol. 10 Preprint at 10.3389/fcell.2022.772230 (2022).

5. Klejchova, M., Silva-Alvim, F. A. L., Blatt, M. R. & Alvim, J. C. Membrane voltage as a dynamic platform for spatiotemporal signaling, physiological, and developmental regulation. Plant Physiology vol. 185 1523–1541 Preprint at 10.1093/plphys/kiab032 (2021).

6. Lobikin, M., Chernet, B., Lobo, D. & Levin, M. Resting potential, oncogene-induced tumorigenesis, and metastasis: The bioelectric basis of cancer in vivo. Phys Biol 9, (2012).

7. Soundarrajan, D. K., Huizar, F. J., Paravitorghabeh, R., Robinett, T. & Zartman, J. J. From spikes to intercellular waves: Tuning intercellular calcium signaling dynamics modulates organ size control. PLoS Comput Biol 17, e1009543 (2021).

8. Srivastava, K. H. et al. Motor control by precisely timed spike patterns. Proceedings of the National Academy of Sciences 114, 1171–1176 (2017).

9. McCaig, C. D., Rajnicek, A. M., Song, B. & Zhao, M. Controlling Cell Behavior Electrically: Current Views and Future Potential. Physiol Rev 85, 943–978 (2005).

10. Levin, M., Thorlin, T., Robinson, K. R., Nogi, T. & Mercola, M. Asymmetries in H/K-ATPase and Cell Membrane Potentials Comprise a Very Early Step in Left-Right Patterning.Cell vol. 111 http://www.cell.com/cgi/ (2002).

11. Blackiston, D. J., McLaughlin, K. A. & Levin, M. Bioelectric controls of cell proliferation: Ion channels, membrane voltage and the cell cycle. Cell Cycle 8, 3527–3536 (2009).

12. Sundelacruz, S., Levin, M. & Kaplan, D. L. Role of Membrane Potential in the Regulation of Cell Proliferation and Differentiation. Stem Cell Rev Rep 5, 231– 246 (2009).

13. Zhao, M. Electrical fields in wound healing—An overriding signal that directs cell migration. Semin Cell Dev Biol 20, 674–682 (2009).

14. Pai, V. P. et al. HCN4 ion channel function is required for early events that regulate anatomical left-right patterning in a nodal and lefty asymmetric gene expression-independent manner. Biol Open 6, 1445–1457 (2017).

15. Beane, W. S., Morokuma, J., Lemire, J. M. & Levin, M. Bioelectric signaling regulates head and organ size during planarian regeneration. Development (Cambridge) 140, 313–322 (2013).

16. Yang, M. & Brackenbury, W. J. Membrane potential and cancer progression. Frontiers in Physiology vol. 4 JUL Preprint at 10.3389/fphys.2013.00185 (2013).

17. Levin, M. Bioelectrical approaches to cancer as a problem of the scaling of the cellular self. Prog Biophys Mol Biol 165, 102–113 (2021).

18. Vaibhav, P. P., Sherry, A., Tal, S., Joan, M. L. & Michael, L. Transmembrane voltage potential controls embryonic eye patterning in Xenopus laevis. Development vol. 139 623 Preprint at 10.1242/dev.077917 (2012).

19. Bates, E. Ion Channels in Development and Cancer. Annu Rev Cell Dev Biol 31, 231–247 (2015).

20. Marchal, G. A. et al. Recent advances and current limitations of available technology to optically manipulate and observe cardiac electrophysiology. Pflugers Arch 475, 1357–1366 (2023).

21. Chorev, E., Epsztein, J., Houweling, A. R., Lee, A. K. & Brecht, M. Electrophysiological recordings from behaving animals—going beyond spikes. Curr Opin Neurobiol 19, 513–519 (2009).

22. Peterka, D. S., Takahashi, H. & Yuste, R. Imaging Voltage in Neurons. Neuron vol. 69 9–21 Preprint at 10.1016/j.neuron.2010.12.010 (2011).

23. Cohen, A. E. & Venkatachalam, V. Bringing bioelectricity to light. Annu Rev Biophys 43, 211–232 (2014).

24. Xu, Y., Zou, P. & Cohen, A. E. Voltage imaging with genetically encoded indicators. Current Opinion in Chemical Biology vol. 39 1–10 Preprint at 10.1016/j.cbpa.2017.04.005 (2017).

25. Pal, A. & Tian, L. Imaging voltage and brain chemistry with genetically encoded sensors and modulators. Current Opinion in Chemical Biology vol. 57 166–176 Preprint at 10.1016/j.cbpa.2020.07.006 (2020).

26. Scott, K. et al. Fast two-photon imaging of subcellular voltage dynamics in neuronal tissue with genetically encoded indicators. doi:10.7554/eLife.25690.001.

27. Bando, Y., Sakamoto, M., Kim, S., Ayzenshtat, I. & Yuste, R. Comparative Evaluation of Genetically Encoded Voltage Indicators. Cell Rep 26, 802–813.e4 (2019).

28. In vivo long-term voltage imaging by genetically encoded voltage indicator reveals spatiotemporal dynamics of neuronal populations during development.

29. Hou, J. H., Kralj, J. M., Douglass, A. D., Engert, F. & Cohen, A. E. Simultaneous mapping of membrane voltage and calcium in zebrafish heart in vivo reveals chamber-specific developmental transitions in ionic currents. Front Physiol 5 **AUG**, (2014).

30. Okumura, K. et al. Optical measurement of neuronal activity in the developing cerebellum of zebrafish using voltage-sensitive dye imaging. Neuroreport 29, 1349–1354 (2018).

31. Levin, M., Pezzulo, G. & Finkelstein, J. M. Endogenous Bioelectric Signaling Networks: Exploiting Voltage Gradients for Control of Growth and Form. (2017) doi:10.1146/annurev-bioeng-071114.

32. Tseng, A. S. & Levin, M. Transducing Bioelectric Signals into Epigenetic Pathways During Tadpole Tail Regeneration. Anatomical Record vol. 295 1541– 1551 Preprint at 10.1002/ar.22495 (2012).

33. Lazzari-Dean, J. R., Gest, A. M. M. & Miller, E. W. Measuring Absolute Membrane Potential Across Space and Time. Annu Rev Biophys 50, 447–468 (2021).

34. Brinks, D., Klein, A. J. & Cohen, A. E. Two-Photon Lifetime Imaging of Voltage Indicating Proteins as a Probe of Absolute Membrane Voltage. Biophys J 109, 914–921 (2015).

35. Lazzari-Dean, J. R., Gest, A. M. & Miller, E. W. Optical estimation of absolute membrane potential using fluorescence lifetime imaging. Elife 8, (2019).

36. Lieschke, G. J. & Currie, P. D. Animal models of human disease: Zebrafish swim into view. Nature Reviews Genetics vol. 8 353–367 Preprint at 10.1038/nrg2091 (2007).

37. Choi, T. Y., Choi, T. I., Lee, Y. R., Choe, S. K. & Kim, C. H. Zebrafish as an animal model for biomedical research. Experimental and Molecular Medicine vol. 53 310–317 Preprint at 10.1038/s12276-021-00571-5 (2021).

38. Shin, J. T. & Fishman, M. C. From zebrafish to human: Modular medical models. Annual Review of Genomics and Human Genetics vol. 3 311–340 Preprint at 10.1146/annurev.genom.3.031402.131506 (2002).

39. White, R. M., et al. Transparent Adult Zebrafish as a Tool for In Vivo Transplantation Analysis. Cell Stem Cell 2, 183–189 (2008).

40. Kibat, C., Krishnan, S., Ramaswamy, M., Baker, B. J. & Jesuthasan, S. Imaging voltage in zebrafish as a route to characterizing a vertebrate functional connectome: promises and pitfalls of genetically encoded indicators. J Neurogenet 30, 80–88 (2016).

41. Böhm, U. L. et al. Voltage imaging identifies spinal circuits that modulate locomotor adaptation in zebrafish. Neuron 110, 1211–1222.e4 (2022).

42. Abdelfattah, A. S. et al. Bright and Photostable Chemigenetic Indicators for Extended in Vivo Voltage Imaging. http://science.sciencemag.org/.

43. Tsutsui, H., Higashijima, S. ichi, Miyawaki, A. & Okamura, Y. Visualizing voltage dynamics in zebrafish heart. Journal of Physiology 588, 2017–2021 (2010).

44. van Opbergen, C. J. M. et al. Optogenetic sensors in the zebrafish heart: A novel in vivo electrophysiological tool to study cardiac arrhythmogenesis. Theranostics 8, 4750–4764 (2018).

45. Tsutsui, H., Karasawa, S., Okamura, Y. & Miyawaki, A. Improving membrane voltage measurements using FRET with new fluorescent proteins. Nat Methods 5, 683–685 (2008).

46. Avitan, L. et al. Spontaneous and evoked activity patterns diverge over development. Elife 10, (2021).

47. Avitan, L. et al. Spontaneous and evoked activity patterns diverge over development. Elife 10, (2021).

48. Gong, Y. et al. High-speed recording of neural spikes in awake mice and flies with a fluorescent voltage sensor. Science (1979) 350, 1361–1366 (2015).

49. Kawakami, K. Tol2: A versatile gene transfer vector in vertebrates. Genome Biology vol. 8 Preprint at 10.1186/gb-2007-8-s1-s7 (2007).

50. Mosimann, C. et al. Ubiquitous transgene expression and Cre-based recombination driven by the ubiquitin promoter in zebrafish. Development 138, 169–177 (2011).

51. Sur, A. et al. Single-cell analysis of shared signatures and transcriptional diversity during zebrafish development. Dev Cell 58, 3028–3047.e12 (2023).

52. Farrell, J. A. et al. Single-cell reconstruction of developmental trajectories during zebrafish embryogenesis. Science (1979) 360, (2018).

53. Brunet, T. et al. The evolutionary origin of bilaterian smooth and striated myocytes. Elife 5, (2016).

54. Kimmel, C. B., Ballard, W. W., Kimmel, S. R., Ullmann, B. & Schilling, T. F. Stages of embryonic development of the zebrafish. Developmental Dynamics 203, 253–310 (1995).

55. Brittijn, S. A. et al. Zebrafish development and regeneration: new tools for biomedical research. Int J Dev Biol 53, 835–850 (2009).

56. Silver, P., Buja, L. & Stull, J. Frequency-dependent myosin light chain phosphorylation in isolated myocardium. J Mol Cell Cardiol 18, 31–37 (1986).

57. Barral, J. M., Hutagalung, A. H., Brinker, A., Hartl, F. U. & Epstein, H. F. Role of the Myosin Assembly Protein UNC-45 as a Molecular Chaperone for Myosin. Science (1979) 295, 669–671 (2002).

58. Rezaei, M. et al. Establishment of a Transgenic Zebrafish Expressing GFP in the Skeletal Muscle as an Ornamental Fish. Galen Medical Journal 8, 1068 (2019).

59. Rudeck, S., et al. A compact unc45b-promoter drives muscle-specific expression in zebrafish and mouse. Genesis vol. 54 431–438 Preprint at 10.1002/dvg.22953 (2016).

60. Etheridge, L., Diiorio, P. & Sagerström, C. G. A zebrafish unc-45–related gene expressed during muscle development. Developmental Dynamics 224, 457– 460 (2002).

61. Wohlgemuth, S. L., Crawford, B. D. & Pilgrim, D. B. The myosin co-chaperone UNC-45 is required for skeletal and cardiac muscle function in zebrafish. Dev Biol 303, 483–492 (2007).

62. Whitesell, T. R. et al. An α-smooth muscle actin (acta2/αsma) zebrafish transgenic line marking vascular mural cells and visceral smooth muscle cells. PLoS One 9, (2014).

63. Kim, C.-H., et al. HIUROSCI[NC[ l[ITlll Zebrafish Elav/HuC Homologue as a Very Early Neuronal Marker. Neuroscience Letters vol. 216 (1996).

64. Liy, J. Early Development of Functional Spatial Maps in the Zebrafish Olfactory Bulb. Journal of Neuroscience 25, 5784–5795 (2005).

65. Fuller, C. L., Yettaw, H. K. & Byrd, C. A. Mitral cells in the olfactory bulb of adult zebrafish (Danio rerio): Morphology and distribution. Journal of Comparative Neurology 499, 218–230 (2006).

66. Sagasti, A., Guido, M. R., Raible, D. W. & Schier, A. F. Repulsive interactions shape the morphologies and functional arrangement of zebrafish peripheral sensory arbors. Current Biology 15, 804–814 (2005).

67. Takahashi, M., Narushima, M. & Oda, Y. In Vivo Imaging of Functional Inhibitory Networks on the Mauthner Cell of Larval Zebrafish. The Journal of Neuroscience 22, 3929–3938 (2002).

68. Juárez-Morales, J. L., Martinez-De Luna, R. I., Zuber, M. E., Roberts, A. & Lewis, K. E. Zebrafish transgenic constructs label specific neurons in Xenopus laevis spinal cord and identify frog V0v spinal neurons. Dev Neurobiol 77, 1007–1020 (2017).

69. Harris, J. M., Yu-Der Wang, A. & Arlotta, P. Optogenetic axon guidance in embryonic zebrafish. STAR Protoc 2, (2021).

70. Mueller, T. & Wullimann, M. F. Expression Domains of NeuroD (Nrd) in the Early Postembryonic Zebrafish Brain. (2002).

71. Khuansuwan, S., Barnhill, L. M., Cheng, S. & Bronstein, J. M. A novel transgenic zebrafish line allows for in vivo quantification of autophagic activity in neurons. Autophagy 15, 1322–1332 (2019).

72. Sato, A. & Takeda, H. Neuronal Subtypes Are Specified by the Level of *neurod* Expression in the Zebrafish Lateral Line. The Journal of Neuroscience 33, 556– 562 (2013).

73. Mueller, T. & Wullimann, M. F. BrdU-, neuroD (nrd)- and Hu-studies reveal unusual non-ventricular neurogenesis in the postembryonic zebrafish forebrain. Mech Dev 117, 123–135 (2002).

74. Hodgkin, A. L. & Huxley, A. F. A quantitative description of membrane current and its application to conduction and excitation in nerve. J Physiol 117, 500– 544 (1952).

75. Zhang, X. M., Yokoyama, T. & Sakamoto, M. Imaging Voltage with Microbial Rhodopsins. Frontiers in Molecular Biosciences vol. 8 Preprint at 10.3389/fmolb.2021.738829 (2021).

76. McLean, D. L. Voltage imaging and spinal circuits get along swimmingly. Neuron vol. 110 1093–1094 Preprint at 10.1016/j.neuron.2022.03.024 (2022).

77. Knöpfel, T. & Song, C. Optical voltage imaging in neurons: moving from technology development to practical tool. Nat Rev Neurosci 20, 719–727 (2019).

78. Boulanger-Weill, J. & Sumbre, G. Functional Integration of Newborn Neurons in the Zebrafish Optic Tectum. Front Cell Dev Biol 7, (2019).

79. Chapouton, P. & Bally-Cuif, L. Neurogenesis. in 163–206 (2004). doi:10.1016/S0091-679X(04)76010-0.

80. Wullimann, M. F. Secondary neurogenesis and telencephalic organization in zebrafish and mice: a brief review. Integr Zool 4, 123–133 (2009).

81. Bakkers, J. Zebrafish as a model to study cardiac development and human cardiac disease. Cardiovasc Res 91, 279–288 (2011).

82. Giardoglou, P. & Beis, D. On Zebrafish Disease Models and Matters of the Heart. Biomedicines 7, 15 (2019).

83. Várkuti, B. H. et al. A highly soluble, non-phototoxic, non-fluorescent blebbistatin derivative. Sci Rep 6, (2016).

84. Chi, N. C. et al. Genetic and physiologic dissection of the vertebrate cardiac conduction system. PLoS Biol 6, 1006–1019 (2008).

85. Nemtsas, P., Wettwer, E., Christ, T., Weidinger, G. & Ravens, U. Adult zebrafish heart as a model for human heart? An electrophysiological study. J Mol Cell Cardiol 48, 161–171 (2010).

86. Silic, M. R. & Zhang, G. Bioelectricity in Developmental Patterning and Size Control: Evidence and Genetically Encoded Tools in the Zebrafish Model. Cells 12, 1148 (2023).

87. Schmidt, R., Strähle, U. & Scholpp, S. Neurogenesis in Zebrafish-from Embryo to Adult. http://www.neuraldevelopment.com/content/8/1/3 (2013).

88. Warp, E. et al. Emergence of patterned activity in the developing zebrafish spinal cord. Current Biology 22, 93–102 (2012).

89. David Wong-Campos, J., et al. Voltage dynamics of dendritic integration and back-propagation in vivo. doi:10.1101/2023.05.25.542363.

90. Ince-Dunn, G. et al. Neuronal Elav-like (Hu) Proteins Regulate RNA Splicing and Abundance to Control Glutamate Levels and Neuronal Excitability. Neuron 75, 1067–1080 (2012).

91. Piatkevich, K. D. et al. Population imaging of neural activity in awake behaving mice. Nature 574, 413–417 (2019).

92. Kim, B. B. et al. A red fluorescent protein with improved monomericity enables ratiometric voltage imaging with ASAP3. Sci Rep 12, 3678 (2022).

93. Biasci, V. et al. Optogenetic manipulation of cardiac electrical dynamics using sub-threshold illumination: dissecting the role of cardiac alternans in terminating rapid rhythms. Basic Res Cardiol 117, 25 (2022).

94. Levin, M. Bioelectric signaling: Reprogrammable circuits underlying embryogenesis, regeneration, and cancer. Cell 184, 1971–1989 (2021).

95. Pai, V. P. et al. HCN2 Rescues brain defects by enforcing endogenous voltage pre-patterns. Nat Commun 9, 998 (2018).

96. Philipp Rühl, A., et al. An ultrasensitive genetically encoded voltage indicator uncovers the electrical activity of non-excitable cells. doi:10.1101/2023.10.05.560122.

97. Pnevmatikakis, E. A. & Giovannucci, A. NoRMCorre: An online algorithm for piecewise rigid motion correction of calcium imaging data. J Neurosci Methods 291, 83–94 (2017).

98. Kleinfeld, D. et al. CaImAn an open source tool for scalable calcium imaging data analysis. (2019) doi:10.7554/eLife.38173.001.

99. Eom, M. et al. Statistically unbiased prediction enables accurate denoising of voltage imaging data. Nat Methods 20, 1581–1592 (2023).

100. Cai, C. et al. VolPy: Automated and scalable analysis pipelines for voltage imaging datasets. PLoS Comput Biol 17, (2021).

